# CytoNormPy enables a fast and scalable removal of batch effects in cytometry datasets

**DOI:** 10.1101/2024.07.19.604225

**Authors:** Tarik Exner, Nicolaj Hackert, Luca Leomazzi, Sofie Van Gassen, Yvan Saeys, Hanns-Martin Lorenz, Ricardo Grieshaber-Bouyer

**Affiliations:** Internal Medicine V, Hematology, Oncology and Rheumatology, Heidelberg University Hospital, Heidelberg, Germany; Institute for Immunology, Heidelberg University Hospital, Germany; Data Mining and Modeling for Biomedicine group, VIB Center for Inflammation Research, Ghent, Belgium; Department of Applied Mathematics, Computer Science and Statistics, Ghent University, Ghent, Belgium; Deutsches Zentrum Immuntherapie (DZI), Friedrich-Alexander-Universität Erlangen-Nürnberg and Universitätsklinikum Erlangen, Erlangen, Germany; Department of Internal Medicine 5 – Hematology and Oncology, Friedrich-Alexander-Universität Erlangen-Nürnberg and University Hospital Erlangen, Erlangen, Germany; Department of Internal Medicine 3 – Rheumatology and Immunology, Friedrich-Alexander-Universität Erlangen-Nürnberg and University Hospital Erlangen, Erlangen, Germany

## Abstract

**Motivation:** We present a python implementation of the widely used CytoNorm algorithm for the removal of batch effects.

**Results:** Our implementation ran up to 85% faster than its R counterpart, while being fully compatible with common single-cell data structures and -frameworks of python. We extend the previous functionality by adding common clustering algorithms and provide key visualizations of the algorithm and its evaluation.

**Availability and implementation:** The CytoNormPy implementation is freely available on GitHub: https://github.com/TarikExner/CytoNormPy.

## Introduction

Cytometry is an essential technique to measure multiple features of single cells. As the scale of cytometry studies grows and longitudinal studies are performed, not all samples can be acquired simultaneously, which introduces batch effects into the data. Therefore, the analysis of cytometry data increasingly necessitates accurate and efficient methods for batch correction.

CytoNorm has recently been described as an algorithm for the removal of batch effects using cluster-specific marker normalization across batches (1). The algorithm uses reference samples measured in all batches to quantify technical variation between different batches in the dataset (summarized in **Supplementary Figure 1**). CytoNorm has been widely adopted in the analysis of cytometry data in fundamental science (e.g. (2), (3)) and is currently the most cited algorithm among peer-reviewed methods for the removal of batch effects in mass cytometry (iMUBAC (4), CytofRUV (5), CytofBatchAdjust (6) and cyCombine (7)).

CytoNorm is currently implemented in R, one of the most widely used programming languages for data analysis, including cytometry. However, python has become increasingly popular in the analysis of single-cell data, as it offers a generalizable and flexible data structure (*anndata* (8)) and accompanying frameworks like the *scverse (9)* ecosystem, *scanpy* (10) and *pytometry* (10, 11), among others. Further, many popular machine/deep learning frameworks provide python APIs and python often outperforms R in terms of scalability and speed in many applications.

Here, we present *cytonormpy*, a python implementation of the CytoNorm algorithm. We show that *cytonormpy* runs significantly faster than CytoNorm, yielding near identical results with the original implementation and offering full compatibility with python single cell analysis data structures. We extend the original capabilities and add suitable clustering algorithms for cytometry data (12-15). Finally, we provide visualization techniques to display key steps of the algorithm and its results.

## Materials and Methods

### Implementation details

The implementation described here follows the steps implemented in the R package. For the use of .fcs files, the reference files from a sample present in all batches are first read using *flowio* (16), and annotated using user-provided metadata while pre-assembled *anndata (8)* objects can directly be passed into the function. Next, the data are optionally transformed using one of the implemented transformation methods in flowutils (16), namely *log, logicle, arcsinh* and *hyperlog* transformation. Data are subsequently clustered using the FlowSOM (17, 18), KMeans, MeanShift or AffinityPropagation (scikit-learn (19) implementation) algorithm. A user-defined number of quantiles (default: 99) is then calculated per marker, cluster, channel and batch, which are saved into a pre-allocated *numpy* nd-array for more efficient data storage and -access. Quantized expression values of markers of cells in clusters with less than a user defined number of cells (default: 50 cells per cluster) are masked out and the corresponding clusters are excluded from further calculations. The desired marker distribution across batches (goal distribution) is calculated by taking the mean of the expression quantiles over batches. Alternatively, the median can be used, or the expression values of a specific batch can be selected. Next, spline functions are calculated for each marker and cluster using the batch-specific quantile expression values and the respective goal distribution.

The fitting of the spline function follows the R implementation exactly. First, duplicate values within the quantile expression values are removed using a custom implementation of R’s *regularize.values* function using the mean as a tie-function. Subsequently, a cubic hermite spline is calculated using the *scipy* (20) implementation. The interpolating tangents are selected using the Fritsch-Carlson method (21) using a custom implementation modeled after the R stats library. The spline function outside of the fitted values is linearly extended using the slope of the first and last datapoint.

The final normalization step of non-reference files is carried out as follow: First, the files are read concurrently using multi-threading. The data are subsequently mapped onto the clusters obtained from the clustering algorithms trained on the reference data. The cluster-specific expression values are finally normalized using the respective spline functions. FCS data are written to the hard drive, while *anndata* objects are returned.

For the evaluation of batch-effect removal, we implemented the calculation of Earth Mover’s distance according to the R implementation of CytoNorm. Finally, in order to quantify the conservation of biological signal, the median absolute deviation per channel in the original and the normalized samples are computed.

For functions carrying out numerical operations, including the regularize_values function, the Fritsch-Carlson tangent selection and quantile calculation, we make use of the *numba (22)* JIT (just in time) compilation package with eager compilation in nopython mode to take advantage of the increased performance of precompiled code in python.

### Dataset preprocessing and benchmarking

For benchmarking and reproducibility comparison, we used three datasets in total (compare **Table 1**; detailed sample-wise descriptions and panel information are provided in the **Supplemental Material**). The Van Gassen dataset (17) consists of two healthy donors with stimulated and unstimulated data measured over 10 batches (40 samples). For the benchmark, samples of healthy control 1 were used as reference, while samples of healthy control 2 were normalized. The dataset assembled by Trussart et al. (5) contained samples of healthy controls (n=3) and patients with chronic leukocytic leukemia (n=3) in two batches. The dataset was first pre-processed as described in their methods section and the sample HC1 and CLL2 were used as a reference. The third dataset (Dana Farber Cancer Institute (7), DFCI) consisted of three batches, of which the third one was used as reference. The data were split into two panels with 15 overlapping markers and both panels were considered a single batch, following the procedure of the original publication (7). The samples HD05 and CLL08 were used as references. For the benchmark, only the channels corresponding to antigens measured by antibodies were selected. All data were *arcsinh* transformed using a cofactor of 5. The reference data were clustered without subsampling using the FlowSOM algorithm with a grid size of 5×5 and a total of 10 clusters and normalization was applied for each cluster as described above. As a control, the same analysis was repeated without clustering in order to remove the effect of inter-language deviation due to small differences in cluster assignment comparing the FlowSOM implementations in R and python. Execution time was measured using the *time* module in python (version 3.10) and the Sys library in R (version 4.4.0), respectively. For the benchmark, CytoNorm version 2.0.2 was used.

**Table 1:** Datasets used in this study. Statistics for the number of batches within the respective dataset (n batches) as well as the number of individual files (n samples), the channel count (n channels) and the number of cells in the reference and validation subsets (n cells) are given. All datasets are mass cytometry datasets.

| Dataset | FlowRepository<br>Accession | n<br>batches | n_samples<br>(reference/validation) | n_channels | n_cells<br>(reference/validation) |
| --- | --- | --- | --- | --- | --- |
| Van<br>Gassen | FR-FCM-Z247 | 10 | 40 (20/20) | 37 | 6.193.423<br>(2.842.208/3.351.215) |
| Trussart | FR-FCM-Z2L2 | 2 | 24 (4/20) | 34 | 8.589.739<br>(2.095.046/6.494.693) |
| DFCI<br>(batch<br>3) | FR-FCM-Z52G | 2 | 39 (4/35) | 15 | 1.910.861<br>(508.055/1.402.806) |

## Data availability

All datasets used in our study are available at FlowRepository. For the accession numbers refer to **Table 1**.

## Code availability

The cytonormpy package is freely available on GitHub: https://github.com/TarikExner/CytoNormPy. The code to reproduce the data and figures used in this paper is provided in the **Supplementary Material**.

## Results and Discussion

### Highly enhanced performance of the python implementation

We first measured the computation time of our implementation in direct comparison to the R implementation on an 8 core machine and 32GB RAM using three datasets (**Table 1** and **Methods**). Using the default implementation including the clustering step, we found that the python implementation led to a speed increase of 72%-85% (**Figure 1A**). Upon further inspection of the runtime of the individual steps, we noticed that most of the runtime was spent in I/O operations and the clustering step (**Supplementary Figure 2A**). As the R implementation uses frequent saving to the hard-drive to save cluster specific data, the observed speed difference is best explained by the increased amount of I/O operations, although it should be mentioned that the FlowSOM clustering itself has a lower runtime in python compared to its R counterpart (18). In order to evaluate the performance independent of the intra-algorithm I/O operations and FlowSOM clustering itself, we compared the runtimes while skipping the clustering step. Here, our implementation showed a noticeable performance benefit of 27%-43% compared to the implementation in R.

**Figure 1:**
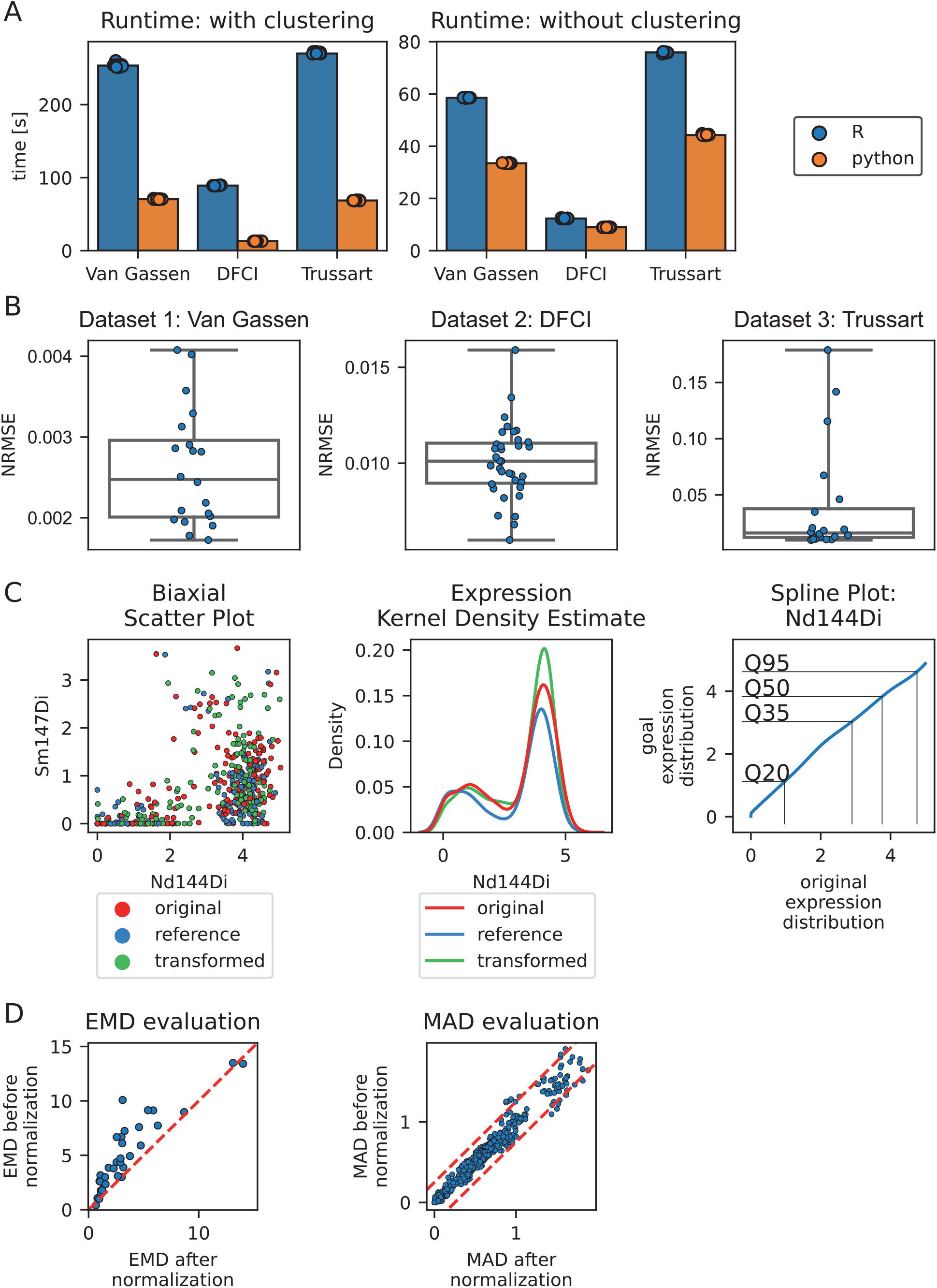
Benchmarking and cross-language comparison. **A** Runtime comparison. Runtime was determined on the indicated datasets for 10 runs. Displayed is the median runtime with (left plot) and without the clustering step (right) showing a significant improvement of runtime in the python implementation compared to R, especially if the default implementation including the clustering step was used. **B** Deviation of normalized expression values comparing python and R. Data were processed as described above, including the clustering step. The root mean squared error (RMSE) between the normalized values calculated from the R implementation and the python implementation was calculated per file and channel, normalized to the respective channel range and plotted as average over all channels per file (represented by the data points). **C** Visualization of the normalization procedure. Cytonormpy implements a scatter plot (left) and a kernel density estimate (middle) in order to visualize the effect of normalization. A splineplot (right) is used to visualize the fitted spline function where individual, user-defined quantile values can be displayed. **D** Visualization of the evaluation metrics. Earth Mover’s distances are computed before and after normalization and are visualized as a scatter plot (left plot). The red dashed line marks a deviation of zero, allowing seamless interpretation regarding the change. Median absolute deviations (MAD) are calculated per file and channel and are plotted as a scatter plot comparing the respective values before and after normalization (right plot). The red dashed line represents a user-defined cutoff for the respective MAD change (here: ±0.25).

### Cytonormpy yields almost identical results compared to CytoNorm

We next tested the difference of the resulting normalized values comparing the R and python implementation. We found that the values were highly concordant as judged by the normalized RMSE values per file (**Figure 1B**). We could trace the numerical differences back to small deviations in cluster assignment of the cells, as skipping the clustering step resulted in identical values for the batch-normalized values between python and R (**Supplementary Figure 2B**).

### Visualization techniques

We implemented a dedicated class for data visualization. In addition to a normal biaxial scatter plot, we provide tools to visualize the normalization impact using kernel density estimates. A splineplot was implemented that plots the expression values and the corresponding goal distribution as a line plot, labelling user defined quantiles for a clear display of the underlying fitted function (**Figure 1C**). For the visualization of the evaluation metrics Earth Mover’s distance (EMD) and the median absolute deviation (MAD), scatter plots are provided that plot the respective metrics before and after normalization (**Figure 1D**). The resulting plots are fully customizable using the widely used matplotlib and seaborn libraries.

## Conclusion

We present a python implementation of the CytoNorm algorithm, which has been widely used to remove batch effects by reference-driven data normalization. While being fully compatible with existing frameworks like *scanpy* or *pytometry*, we show that our implementation outperforms the R implementation significantly in terms of runtime while yielding highly comparable results. Finally, dedicated visualization capabilities enable a comprehensive diagnostic understanding of the algorithm by users and allow to quantify the effect of batch-normalization and conservation of biological signal.

## Acknowledgements

T.E. was supported by an MD/PhD fellowship from the Medical Faculty of Heidelberg.

L.L. has received funding from the European Union’s Horizon 2021 research and innovation programme under the Marie Skłodowska-Curie grant agreement No 101072891. S.V.G. is supported by an FWO postdoctoral research grant (1272823N, Research Foundation – Flanders). This work was supported by grants to R.G.-B. from Deutsche Forschungsgemeinschaft (DFG, GR 5979/2-1, 517717827), Else Kröner-Fresenius-Stiftung (2022_EKEA.72), the IZKF Erlangen (grant N10), state of Baden-Wuerttemberg within the Centers for Personalized Medicine Baden-Wuerttemberg (ZPM) and a research grant from the German Society for Rheumatology (DGRh). The authors acknowledge support by the state of Baden-Württemberg through bwHPC and the German Research Foundation (DFG) through grant INST 35/1597-1 FUGG.

---

**Supplementary Figure S1:**
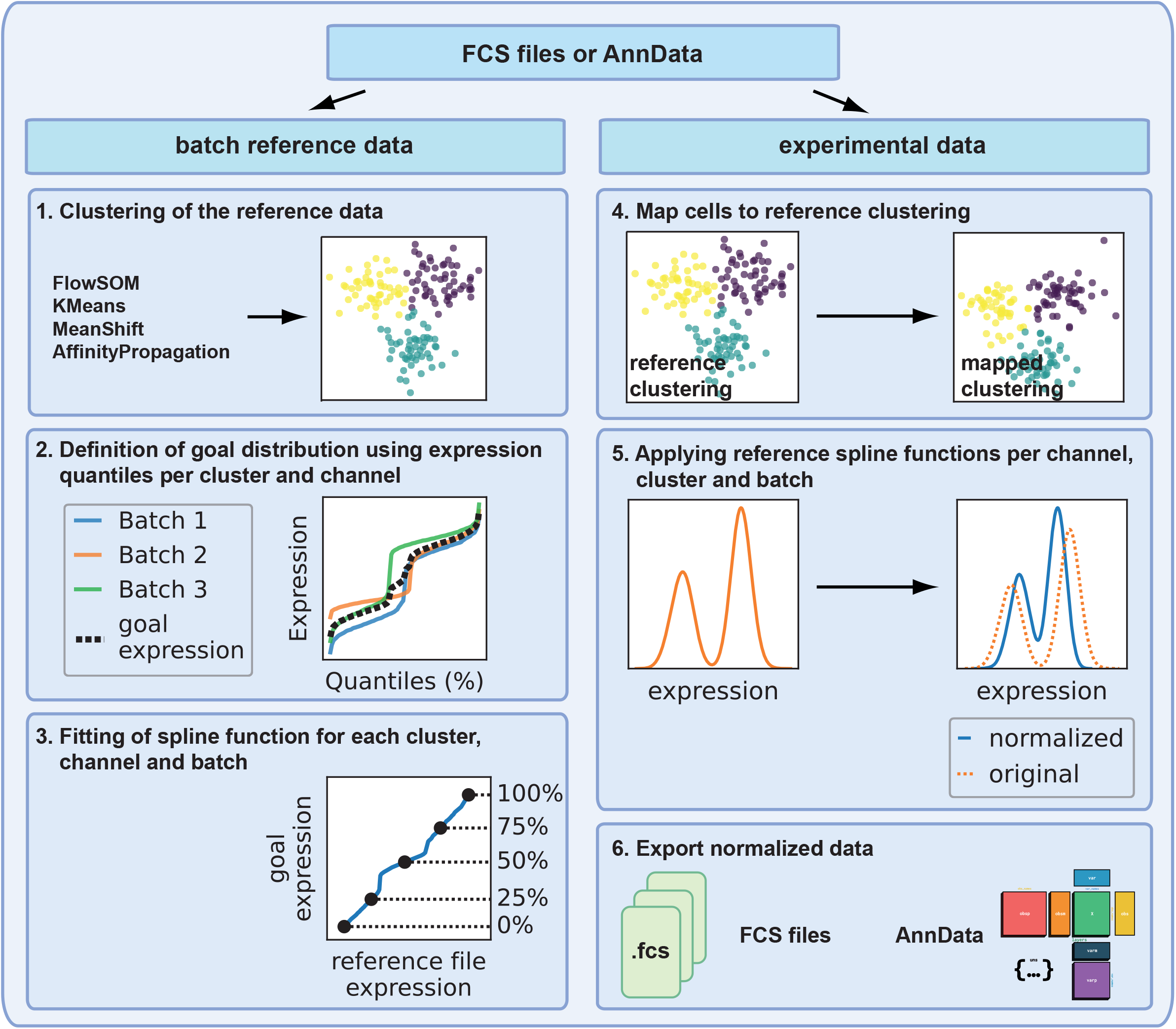
CytoNorm/CytoNormPy algorithm overview. FCS files or anndata objects are divided into reference data (left side) and experimental data (right side) where reference data are comprised of samples that have been measured in all batches. Reference data are subsequently clustered (1) using one of the implemented clustering algorithms. Expression values are quantized, and a desired goal expression is defined, which defaults to the mean of all quantized expression values over the batches (2). A spline function between the expression values and the goal expression is fitted for each channel, cluster, and batch (3). The experimental data are then mapped to the reference cluster space (4) and the corresponding spline functions are used to transform the data (5). Finally, the data are either exported as .fcs files or returned as an anndata object containing the normalized expression values (6).

**Supplementary Figure S2:**
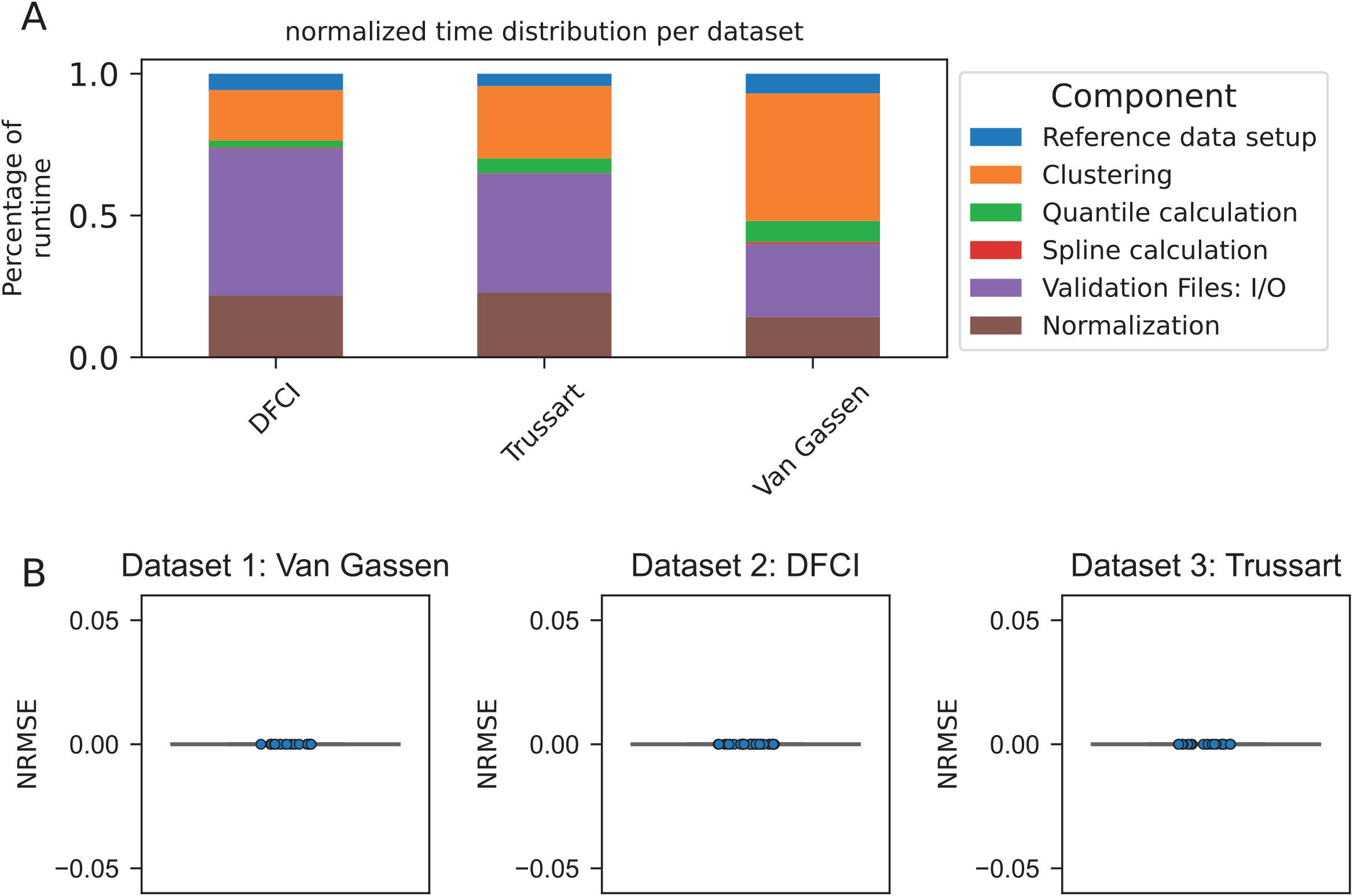
Function benchmark and comparison. **A** Runtime comparison of the individual function parts. The runtime was measured for the indicated steps of cytonormpy. The majority of runtime is spent in read/write (I/O) and clustering operations. The relative difference between the ratios is best explained by the number of cells in the reference- and validation-set, respectively (compare Table 1). **B** Deviation of normalized expression values. Data were subjected to the cytonormpy algorithm, omitting the clustering step. The root mean squared error (RMSE) between the normalized values calculated from the R implementation and the python implementation was calculated per file and channel, normalized to the respective channel range and plotted as average over all channels per file (represented by the data points).

