## Supplementary material for "CytoNormPy enables a fast and scalable removal of batch effects in cytometry datasets": cytonorm_python.py

```
import cytonormpy as cnp
from cytonormpy import FCSFile
import os

import pandas as pd
import anndata as ad
import numpy as np

from pathlib import Path
from sklearn.metrics import root_mean_squared_error
import gc
from time import time

DATASET_INPUT_DIRS: dict[str, os.PathLike] = {
    "van_gassen": Path("../datasets/van_gassen_2/data/"),
    "dfci": Path("../datasets/DFCIb3/data/"),
    "trussard": Path("../datasets/Trussard/data/")
}

PYTHON_INPUT_DIRS: dict[str, os.PathLike] = {
    "van_gassen_clustered": Path("../datasets/van_gassen_2/data/normalized/clustered/python/"),
    "van_gassen_unclustered": Path("../datasets/van_gassen_2/data/normalized/unclustered/python/"),
    "dfci_clustered": Path("../datasets/DFCIb3/data/normalized/clustered/python/"),
    "dfci_unclustered": Path("../datasets/DFCIb3/data/normalized/unclustered/python/"),
    "trussard_clustered": Path("../datasets/Trussard/data/normalized/clustered/python/"),
    "trussard_unclustered": Path("../datasets/Trussard/data/normalized/unclustered/python/"),
}

R_INPUT_DIRS: dict[str, os.PathLike] = {
    "van_gassen_clustered": Path("../datasets/van_gassen_2/data/normalized/clustered/R/"),
    "van_gassen_unclustered": Path("../datasets/van_gassen_2/data/normalized/unclustered/R/"),
    "dfci_clustered": Path("../datasets/DFCIb3/data/normalized/clustered/R/"),
    "dfci_unclustered": Path("../datasets/DFCIb3/data/normalized/unclustered/R/"),
    "trussard_clustered": Path("../datasets/Trussard/data/normalized/clustered/R/"),
    "trussard_unclustered": Path("../datasets/Trussard/data/normalized/unclustered/R/"),
}

OUTPUT_DIRS: dict[str, os.PathLike] = {
    "clustered": Path("normalized/clustered/python/"),
    "unclustered": Path("normalized/unclustered/python/")
}

DATASETS = ["van_gassen", "dfci", "trussard"]

def _write_benchmark(line: str,
                     init: bool = False,
                     functionwise: bool = False) -> None:
    line += "\n"
    file_name = "python_benchmark.out" if not functionwise else "python_benchmark_functionwise.out"
    if init:
        with open(file_name, "w") as output_file:
            output_file.writelines(line)
    else:
        with open(file_name, "a") as output_file:
            output_file.writelines(line)
    return

def _get_coding_detectors(input_dir) -> list[str]:
    return pd.read_csv(os.path.join(input_dir, "coding_detectors.txt"), header = None)[0].tolist()

def _results_file_name(dataset: str, clustered: str, original_comp: bool = False) -> str:
    if original_comp:
        return f"{dataset}_{clustered}_normalization.csv"
    return f"{dataset}_{clustered}.csv"

def _run_cytonorm_function_by_function(input_dir: os.PathLike,
                                       output_dir: os.PathLike,
                                       clustered: bool) -> str:

    coding_detectors = _get_coding_detectors(input_dir)

    cn = cnp.CytoNorm()

    t = cnp.AsinhTransformer()
    cn.add_transformer(t)

    if clustered:
        fs = cnp.FlowSOM(n_clusters = 10,
                         xdim = 5,
                         ydim = 5)
        cn.add_clusterer(fs)


    metadata = pd.read_csv(os.path.join(input_dir, "metadata.csv"), sep = ";")
    time_string = ""
    start = time()
    cn.run_fcs_data_setup(input_directory = input_dir,
                          output_directory = output_dir,
                          channels = coding_detectors,
                          metadata = metadata,
                          truncate_max_range = False)
    time_string += str(time()-start)

    if clustered:
        start = time()
        cn.run_clustering(cluster_cv_threshold = 2)
        time_string += "," + str(time()-start)
    else:
        time_string += "," + str(-1)

    start = time()
    cn.calculate_quantiles()
    time_string += "," + str(time()-start)

    start = time()
    cn.calculate_splines(goal = "batch_mean")
    time_string += "," + str(time()-start)

    # we transform "manually" in order to better visualize the difference I/O and transformation

    # cn.transform_data()
    io_read = []
    transform = []
    io_write = []

    for file in cn._datahandler.validation_file_names:
        start = time()
        df = cn._datahandler.get_dataframe(file_name = file)
        io_read.append(time()-start)

        batch = cn._datahandler.get_batch(file_name = file)

        start = time()
        df = cn._normalize_file(df = df,
                                batch = batch)
        transform.append(time()-start)

        start = time()
        cn._datahandler.write(file_name = file,
                              data = df)
        io_write.append(time()-start)

    io_read = np.sum(io_read)
    transform = np.sum(transform)
    io_write = np.sum(io_write)

    time_string += "," + str(io_read)
    time_string += "," + str(transform)
    time_string += "," + str(io_write)

    return time_string


def _run_cytonorm(input_dir: os.PathLike,
                  output_dir: os.PathLike,
                  clustered: bool) -> None:

    coding_detectors = _get_coding_detectors(input_dir)

    cn = cnp.CytoNorm()

    t = cnp.AsinhTransformer()
    cn.add_transformer(t)

    if clustered:
        fs = cnp.FlowSOM(n_clusters = 10,
                         xdim = 5,
                         ydim = 5)
        cn.add_clusterer(fs)

    metadata = pd.read_csv(os.path.join(input_dir, "metadata.csv"), sep = ";")
    cn.run_fcs_data_setup(input_directory = input_dir,
                          output_directory = output_dir,
                          channels = coding_detectors,
                          metadata = metadata,
                          truncate_max_range = False)

    if clustered:
        cn.run_clustering(cluster_cv_threshold = 2)

    cn.calculate_quantiles()
    cn.calculate_splines(goal = "batch_mean")
    cn.normalize_data()

    return


def _create_output_dir_string(input_dir,
                              clustering: str) -> os.PathLike:
    return Path(os.path.join(input_dir, OUTPUT_DIRS[clustering]))


def _create_anndata(input_dir: os.PathLike,
                    metadata_dir: os.PathLike,
                    original: bool = False) -> ad.AnnData:
    adatas = []
    metadata = pd.read_csv(metadata_dir, sep = ";")
    if not original:
        metadata["file_name"] = [f"Norm_{filename}" for filename in metadata["file_name"].tolist()]
    ref_file_names = metadata.loc[metadata["reference"] == "other", "file_name"].tolist()
    for file in ref_file_names:
        fcs = FCSFile(
            input_directory = input_dir,
            file_name = file,
            truncate_max_range = False
        )
        events = fcs.original_events
        md_row = metadata.loc[
            metadata["file_name"] == file, :
        ].to_numpy()
        obs = np.repeat(
            md_row,
            events.shape[0],
            axis = 0
        )
        var_frame = fcs.channels
        obs_frame = pd.DataFrame(
            data = obs,
            columns = metadata.columns,
            index = pd.Index([str(i) for i in range(events.shape[0])])
        )
        adata = ad.AnnData(
            obs = obs_frame,
            var = var_frame,
            layers = {"compensated": events}
        )
        adata.obs_names_make_unique()
        adata.var_names_make_unique()
        adatas.append(adata)
    
    dataset = ad.concat(adatas, axis = 0, join = "outer", merge = "same")
    dataset.obs = dataset.obs.astype(str)
    dataset.var = dataset.var.astype(str)
    dataset.obs_names_make_unique()
    dataset.var_names_make_unique()
    return dataset

def _clip_df(df) -> pd.DataFrame:
    return df.clip(
        lower = np.quantile(df, 0, axis = 0),
        upper = np.quantile(df, 0.999, axis = 0)
    )

def _select_file_name_and_detectors_and_clip(df, file_name, coding_detectors) -> pd.DataFrame:
    return _clip_df(df.loc[df["file_name"] == file_name, coding_detectors])

def _calculate_NRMSE(df1: pd.DataFrame,
                     df2: pd.DataFrame) -> float:
    interquartile_ranges = df2.quantile(0.999) - df2.quantile(0)
    nrmse = pd.DataFrame(columns = df1.columns,
                         index = ["NRMSE"])
    for det in df1.columns:
        nrmse.loc[nrmse.index == "NRMSE", det] = root_mean_squared_error(
            df1[det],
            df2[det]
        )
    return np.average(
        nrmse / interquartile_ranges
    )

def _calculate_differences(orig_dir: os.PathLike,
                           python_dir: os.PathLike,
                           r_dir: os.PathLike,
                           dataset: str,
                           clustered: str) -> None:

    coding_detectors = _get_coding_detectors(orig_dir)
    metadata_dir = Path(os.path.join(orig_dir, "metadata.csv"))

    original = _create_anndata(
        orig_dir,
        metadata_dir,
        original = True
    )
    python = _create_anndata(
        python_dir,
        metadata_dir,
    )
    r = _create_anndata(
        r_dir,
        metadata_dir,
    )
    original_df = original.to_df(layer = "compensated")
    python_df = python.to_df(layer = "compensated")
    r_df = r.to_df(layer = "compensated")
    
    assert python_df["Time"].tolist() == r_df["Time"].tolist()
    
    python_df[python.obs.columns] = python.obs
    r_df[r.obs.columns] = r.obs
    original_df[original.obs.columns] = original.obs


    res = pd.DataFrame(columns = ["NRMSE"], index = python_df["file_name"].unique())

    for file in python_df["file_name"].unique():
        res.loc[res.index == file, "NRMSE"] = _calculate_NRMSE(
            _select_file_name_and_detectors_and_clip(python_df, file, coding_detectors),
            _select_file_name_and_detectors_and_clip(r_df, file, coding_detectors),
        )
    file_name = _results_file_name(dataset, clustered)
    res.to_csv(file_name)

    print("NRMSE saved successfully!")
    gc.collect()  


def run_normalization():
    for dataset in DATASETS:
        for clustering in ["clustered", "unclustered"]:
            input_dir = DATASET_INPUT_DIRS[dataset]
            _run_cytonorm(
                input_dir = input_dir,
                output_dir = _create_output_dir_string(input_dir, clustering),
                clustered = True if clustering == "clustered" else False
            )

def run_comparison() -> None:
    for dataset in DATASETS:
        for clustering in ["clustered", "unclustered"]:
            orig_dir = DATASET_INPUT_DIRS[dataset]
            python_dir = PYTHON_INPUT_DIRS[f"{dataset}_{clustering}"]
            r_dir = R_INPUT_DIRS[f"{dataset}_{clustering}"]
            _calculate_differences(
                orig_dir = orig_dir,
                python_dir = python_dir,
                r_dir = r_dir,
                dataset = dataset,
                clustered = clustering
            )


def run_benchmark():
    _write_benchmark("dataset,clustering,time,iteration", init = True)
    for iteration in range(10):
        for dataset in DATASETS:
            for clustering in ["clustered", "unclustered"]:
                input_dir = DATASET_INPUT_DIRS[dataset]
                start = time()
                _run_cytonorm(
                    input_dir = input_dir,
                    output_dir = _create_output_dir_string(input_dir, clustering),
                    clustered = True if clustering == "clustered" else False
                )
                stop = time()-start
                _write_benchmark(
                    f"{dataset},{'TRUE' if clustering == 'clustered' else 'FALSE'},{stop},{iteration}"
                )
                gc.collect()


def run_benchmark_functionwise():
    _write_benchmark(
        "dataset,clustering,time,iteration,fcs_data_setup,clustering,quantile_calc,spline_calc,val_read,transform,val_write",
        init = True,
        functionwise = True
    )
    for iteration in range(10):
        for dataset in DATASETS:
            for clustering in ["clustered", "unclustered"]:
                input_dir = DATASET_INPUT_DIRS[dataset]
                start = time()
                time_string = _run_cytonorm_function_by_function(
                    input_dir = input_dir,
                    output_dir = _create_output_dir_string(input_dir, clustering),
                    clustered = True if clustering == "clustered" else False
                )
                stop = time()-start
                _write_benchmark(
                    f"{dataset},{'TRUE' if clustering == 'clustered' else 'FALSE'},{stop},{iteration},{time_string}",
                    functionwise = True
                )
                gc.collect()


if __name__ == "__main__":
    run_benchmark()
    print("Benchmark complete!")
    run_benchmark_functionwise()
    print("Functionwise benchmark complete!")
    run_normalization()
    print("Normalization complete!")
    run_comparison()
    print("Comparison complete!")
```
