## Supplementary material for "CytoNormPy enables a fast and scalable removal of batch effects in cytometry datasets": cytonorm_r.R

```
library(dplyr)
library(CytoNorm)
library(flowCore)
library(dplyr)
base_directory <- "../datasets/"

script_dir <- ""

setwd(script_dir)

run_flowsom_clustered <- function(train, validation, channels, transformList, transformList.reverse, input_dir){
  model <- CytoNorm.train(files = train$Path,
                          labels = train$batch,
                          channels = channels,
                          transformList = transformList,
                          FlowSOM.params = list(xdim = 5,
                                                ydim = 5,
                                                nClus = 10,
                                                nCells = 1000000,
                                                scale = FALSE),
                          normMethod.train = QuantileNorm.train,
                          normParams = list(nQ = 99,
                                            goal = "mean"),
                          seed = 1,
                          recalculate = TRUE,
                          verbose = TRUE)
  
  CytoNorm.normalize(model = model,
                     files = validation$Path,
                     labels = validation$batch,
                     transformList = transformList,
                     transformList.reverse = transformList.reverse,
                     normMethod.normalize = QuantileNorm.normalize,
                     outputDir = paste0(input_dir, "normalized/clustered/R/"),
                     prefix = "Norm_",
                     clean = TRUE,
                     verbose = TRUE)
  
}

run_flowsom_unclustered <- function(train, validation, channels, transformList, transformList.reverse, input_dir){
  qnorm <- QuantileNorm.train(files = train$Path,
                              labels = train$batch,
                              channels = channels,
                              transformList = transformList,
                              nQ = 99,
                              goal = "mean")
  QuantileNorm.normalize(model = qnorm,
                         files = validation$Path,
                         labels = validation$batch,
                         transformList = transformList,
                         transformList.reverse = transformList.reverse,
                         outputDir = paste0(input_dir, "normalized/unclustered/R/"),
                         prefix = "Norm_",)
}

run_flowsom <- function(input_dir, clustered){
  
  channels <- read.delim(paste0(input_dir, "coding_detectors.txt"), sep = "\n", header = FALSE)
  channels <- unlist(channels) %>% as.character()
  files <- list.files(input_dir, pattern = "fcs$")
  metadata <- read.csv(paste0(input_dir, "metadata.csv"), sep = ";")
  metadata$Path <- paste0(input_dir, metadata$file_name)
  
  train <- dplyr::filter(metadata, reference == "ref")
  validation <- dplyr::filter(metadata, reference == "other")
  
  transformList <- flowCore::transformList(channels, cytofTransform)
  transformList.reverse <- flowCore::transformList(channels, cytofTransform.reverse)
  
  if(clustered==TRUE){
    run_flowsom_clustered(train, validation, channels, transformList, transformList.reverse, input_dir)
  } else {
    run_flowsom_unclustered(train, validation, channels, transformList, transformList.reverse, input_dir)
  }
  
}

run_normalization <- function(input_dir){
  run_flowsom(input_dir,
              clustered = TRUE)
  run_flowsom(input_dir,
              clustered = FALSE)  
}

# BENCHMARK
datasets <- c("van_gassen", "dfci", "Trussard")
cat("dataset,clustering,time,iteration\n", file = "R_benchmark.out", append = F)
for(iteration in seq(0,9)){
  for(dataset in datasets){
    if(dataset=="van_gassen"){
      input_dir <- paste0(base_directory, "van_gassen_2/data/")
    }
    if(dataset=="Trussard"){
      input_dir <- paste0(base_directory, "Trussard/data/")
    }
    if(dataset=="dfci"){
      input_dir <- paste0(base_directory, "DFCIb3/data/")
    }
    for(clustering in c(TRUE, FALSE)){
      if (dir.exists("tmp")) {
        unlink("tmp", recursive = TRUE)
      }
      start <- Sys.time()
      run_flowsom(input_dir, clustering)
      stop <- Sys.time()
      time_spent <- difftime(stop, start, units = "secs")
      time_spent <- time_spent %>% as.numeric()
      cat(
        paste(
          dataset, clustering, time_spent, paste0(iteration, "\n"),
          sep = ","
        ),
        file = "R_benchmark.out",
        append = TRUE
      )
    }
  }  
}

gc()

# BENCHMARK with RProf
datasets <- c("van_gassen", "dfci", "Trussard")
output_file <- "profiler.out"
cat("dataset,clustering,time,iteration\n", file = "R_benchmark_rprof_int1.out", append = F)
for(iteration in seq(0,9)){
  for(dataset in datasets){
    if(dataset=="van_gassen"){
      input_dir <- paste0(base_directory, "van_gassen_2/data/")
    }
    if(dataset=="Trussard"){
      input_dir <- paste0(base_directory, "Trussard/data/")
    }
    if(dataset=="dfci"){
      input_dir <- paste0(base_directory, "DFCIb3/data/")
    }
    for(clustering in c(TRUE, FALSE)){
      if (dir.exists("tmp")) {
        unlink("tmp", recursive = TRUE)
      }
      Rprof(output_file, interval = 1)
      start <- Sys.time()
      run_flowsom(input_dir, clustering)
      Rprof(NULL)
      res <- summaryRprof(output_file)
      time <- res$sampling.time

      cat(
        paste(
          dataset, clustering, time, paste0(iteration, "\n"),
          sep = ","
        ),
        file = "R_benchmark_rprof_int1.out",
        append = TRUE
      )
    }
  }  
}
 
if (dir.exists("tmp")) {
  unlink("tmp", recursive = TRUE)
}

# DATASET 1 - van Gassen
input_dir <- paste0(base_directory, "van_gassen_2/data/")
run_normalization(input_dir)

if (dir.exists("tmp")) {
  unlink("tmp", recursive = TRUE)
}

# DATASET 2 - DFCIb3
input_dir <- paste0(base_directory, "DFCIb3/data/")
run_normalization(input_dir)

if (dir.exists("tmp")) {
  unlink("tmp", recursive = TRUE)
}

# DATASET 3 - Trussard
input_dir <- paste0(base_directory, "Trussard/data/")
run_normalization(input_dir)

if (dir.exists("tmp")) {
  unlink("tmp", recursive = TRUE)
}

# DATASET 4 - CytoNormRepo
input_dir <- paste0(base_directory, "cytonorm_repo/data/")
run_normalization(input_dir)

if (dir.exists("tmp")) {
  unlink("tmp", recursive = TRUE)
}
```
