## Supplementary material for "CytoNormPy enables a fast and scalable removal of batch effects in cytometry datasets": figure_generation.ipynb

```
# %% [code]
import os

import numpy as np
import pandas as pd
from matplotlib import pyplot as plt
import seaborn as sns

import matplotlib.patches as mpatches
from matplotlib.colors import LogNorm, SymLogNorm

from matplotlib.axes import Axes
from matplotlib.figure import Figure
from matplotlib.gridspec import GridSpec

# %% [code]
import warnings
warnings.filterwarnings(
    action='ignore',
    category=FutureWarning
)
warnings.filterwarnings(
    action='ignore',
    category=UserWarning
)

# %% [code]
benchmark_python = pd.read_csv("python_benchmark.out")
benchmark_python["language"] = "python"

benchmark_r = pd.read_csv("r_benchmark.out")
benchmark_r["language"] = "R"

benchmark = pd.concat([benchmark_r, benchmark_python], axis = 0)
benchmark["dataset"] = [ds.lower() for ds in benchmark["dataset"].tolist()]
benchmark.head()

# %% [code]
median_times = benchmark.groupby(['dataset', 'clustering', 'language'])['time'].median().reset_index()

pivot_table = median_times.pivot_table(index=['dataset', 'clustering'], columns='language', values='time', fill_value=pd.NA)

pivot_table['R/Python'] = (pivot_table['R'] / pivot_table['python']).fillna('N/A')

pivot_table['Python_Reduction_%'] = ((pivot_table['R'] - pivot_table['python']) / pivot_table['R'] * 100).fillna('N/A')

result = pivot_table.reset_index()
result

# %% [code]
DPI = 300

FIGURE_WIDTH_FULL = 6.75
FIGURE_WIDTH_HALF = FIGURE_WIDTH_FULL / 2

FIGURE_HEIGHT_FULL = 9.375
FIGURE_HEIGHT_HALF = FIGURE_HEIGHT_FULL / 2

# %% [code]
def _figure_label(ax: Axes, label, x = 0, y = 1):
    """labels individual subfigures. Requires subgrid to not use figure axis coordinates."""
    ax.text(x,y,label, fontsize = 12)
    return

# %% [code]
def _run_cytonorm(input_dir: str,
                  coding_detectors: list[str],
                  clustered: bool):
    import cytonormpy as cnp
    if os.path.isfile(os.path.join(input_dir, "cytonorm.model")):
        return cnp.read_model(os.path.join(input_dir, "cytonorm.model"))
    output_dir = input_dir + "output"
    cn = cnp.CytoNorm()
    
    t = cnp.AsinhTransformer()
    cn.add_transformer(t)

    if clustered:
        fs = cnp.FlowSOM(n_clusters = 10,
                         xdim = 5,
                         ydim = 5)
        cn.add_clusterer(fs)
    
    metadata = pd.read_csv(os.path.join(input_dir, "metadata.csv"), sep = ";")
    cn.run_fcs_data_setup(input_directory = input_dir,
                          channels = coding_detectors,
                          output_directory = output_dir,
                          metadata = metadata)
    if clustered:
        cn.run_clustering(cluster_cv_threshold = 2)
    
    cn.calculate_quantiles()
    cn.calculate_splines(goal = "batch_mean")
    cn.normalize_data()
    cn.calculate_mad(groupby = ["file_name"])
    cn.calculate_emd()

    cn.save_model(os.path.join(input_dir + "cytonorm.model"))
    

    return cn
    
input_dir = "../datasets/van_gassen_2/data/"
coding_detectors = pd.read_csv(input_dir + "coding_detectors.txt", header = None)[0].tolist()
cn = _run_cytonorm(input_dir, coding_detectors, False)

# %% [code]
def _comp_plot(df: pd.DataFrame,
               axis: Axes,
               title: str):

    plot_params = {
        "data": df,
        "y": "NRMSE",
        "ax": axis
    }
    sns.stripplot(**plot_params,
                  s = 3,
                  linewidth = 0.5,
                  edgecolor = "black")
    sns.boxplot(**plot_params,
                boxprops = dict(facecolor = "white"),
                whis = (0,100))
    axis.set_title(title, fontname = "Arial", fontsize = 10)
    
    axis.set_ylabel(axis.get_ylabel(), fontsize = 8)
    axis.tick_params(bottom = False)
    axis.set_xlabel("")
    if df.max(axis = None) < 0.001:
        axis.set_ylim(-0.06, 0.06)
    axis.set_yticklabels(axis.get_yticklabels(), fontname = "Arial", fontsize = 8)
    return

# %% [code]
def _generate_subfigure_a(fig: Figure,
                          ax: Axes,
                          gs: GridSpec,
                          subfigure_label: str):
    DATASET_MAP = {
        "trussard": "Trussart",
        "dfci": "DFCI",
        "van_gassen": "Van Gassen"
    }

    ax.axis("off")
    _figure_label(ax, subfigure_label, x = -0.1, y = 1.1)
    axes = gs.subgridspec(1,3, wspace = 0.2, width_ratios = [1,1,0.5])

    left_plot = fig.add_subplot(axes[0])
    right_plot = fig.add_subplot(axes[1])
    legend = fig.add_subplot(axes[2])
    
    benchmark_python = pd.read_csv("python_benchmark.out")
    benchmark_python["language"] = "python"
    
    benchmark_r = pd.read_csv("r_benchmark.out")
    benchmark_r["language"] = "R"
    
    benchmark = pd.concat([benchmark_r, benchmark_python], axis = 0)
    benchmark["dataset"] = [ds.lower() for ds in benchmark["dataset"].tolist()]
    benchmark["dataset"] = benchmark["dataset"].map(DATASET_MAP)
    
    
    
    plot_params_clustered = {
        "data": benchmark[benchmark["clustering"] == True],
        "x": "dataset",
        "y": "time",
        "hue": "language",
        "ax": left_plot
    }
    plot_params_unclustered = {
        "data": benchmark[benchmark["clustering"] == False],
        "x": "dataset",
        "y": "time",
        "hue": "language",
        "ax": right_plot
    }
    
    STRIPPLOT_PARAMS = dict(
        dodge = True,
        edgecolor = "black",
        linewidth = 1,
        s = 5
    )
    BARPLOT_PARAMS = dict(
        estimator = "median",
        errorbar = None,
        #facecolor = "white",
        edgecolor = "black",
        linewidth = 1,
        width = 0.8
    )
        
    
    sns.barplot(**plot_params_clustered,
                **BARPLOT_PARAMS)
    sns.stripplot(**plot_params_clustered,
                  **STRIPPLOT_PARAMS)
    
    
    sns.barplot(**plot_params_unclustered,
                **BARPLOT_PARAMS)
    sns.stripplot(**plot_params_unclustered,
                  **STRIPPLOT_PARAMS)

    left_plot.legend().remove()
    
    
    handles, labels = right_plot.get_legend_handles_labels()
    labels = ["R", "python"]
    for ha in handles[:2]:
        ha.set_edgecolor("black")
    legend.axis("off")
    legend.legend(handles, labels,
                      bbox_to_anchor = (-0, 0.5),
                      loc = "center left",
                      fontsize = 8,
                      edgecolor = "black")
    right_plot.legend().remove()
    
    for i, axis in enumerate([left_plot, right_plot]):
        axis.set_xticklabels(axis.get_xticklabels(), rotation = 0, ha = "center", fontsize = 8)
        axis.set_xlabel("")
        axis.tick_params(labelsize = 8)
    right_plot.set_ylabel("")
    left_plot.set_ylabel("time [s]", fontsize = 8)
    left_plot.set_title("Runtime: with clustering", fontsize = 10)
    right_plot.set_title("Runtime: without clustering", fontsize = 10)

# %% [code]
def _generate_subfigure_b(fig: Figure,
                          ax: Axes,
                          gs: GridSpec,
                          subfigure_label: str):

    ax.axis("off")
    _figure_label(ax, subfigure_label, x = -0.1, y = 1.12)
    axes = gs.subgridspec(1,3, wspace = 0.5)

    top_plot = fig.add_subplot(axes[0])
    mid_plot = fig.add_subplot(axes[1])
    bottom_plot = fig.add_subplot(axes[2])
    
    df = pd.read_csv("van_gassen_clustered.csv", index_col = [0])
    _comp_plot(df, top_plot, "Dataset 1: Van Gassen")

    df = pd.read_csv("dfci_clustered.csv", index_col = [0])
    _comp_plot(df, mid_plot, "Dataset 2: DFCI")

    df = pd.read_csv("trussard_clustered.csv", index_col = [0])
    _comp_plot(df, bottom_plot, "Dataset 3: Trussart")

# %% [code]
def _generate_subfigure_c(fig: Figure,
                          ax: Axes,
                          gs: GridSpec,
                          subfigure_label: str):
    ax.axis("off")
    _figure_label(ax, subfigure_label, x = -0.1, y = 1.12)
    axes = gs.subgridspec(2,3, wspace = 0.6, hspace = 0.5, height_ratios = [1, 0.3])
    
    from cytonormpy import Plotter

    left_plot = fig.add_subplot(axes[0,0])

    cnpl = Plotter(cytonorm = cn)
    left_plot = cnpl.scatter(cn._datahandler.validation_file_names[0],
                             x_channel = "Nd144Di",
                             y_channel = "Sm147Di",
                             x_scale = "linear",
                             y_scale = "linear",
                             s = 5,
                             subsample = 500,
                             linewidth = 0.3,
                             edgecolor = "black",
                             palette = "Set1",
                             ax = left_plot,
                             show = False)
    left_plot.set_title("Biaxial\nScatter Plot", fontsize = 10)
    handles, labels = left_plot.get_legend_handles_labels()
    left_plot.legend().remove()
    left_plot_legend = fig.add_subplot(axes[1,0])
    left_plot_legend.axis("off")
    left_plot_legend.legend(handles,
                             labels,
                            bbox_to_anchor = (0, 0.3),
                             loc = "center left",
                             fontsize = 8)


    mid_plot = fig.add_subplot(axes[0,1])
    mid_plot = cnpl.histogram(cn._datahandler.validation_file_names[0],
                               x_channel = "Nd144Di",
                               x_scale = "linear",
                               y_scale = "linear",
                               ax = mid_plot,
                               subsample = 10_000,
                               palette = "Set1",
                               show = False)
    mid_plot.set_title("Expression\nKernel Density Estimate", fontsize = 10)
    legend = mid_plot.legend_
    handles = legend.legendHandles
    labels = [t.get_text() for t in legend.get_texts()]
    mid_plot.legend_.remove()
    mid_plot_legend = fig.add_subplot(axes[1,1])
    mid_plot_legend.axis("off")
    mid_plot_legend.legend(handles,
                             labels,
                            bbox_to_anchor = (0, 0.3),
                             loc = "center left",
                             fontsize = 8)
    right_plot = fig.add_subplot(axes[0,2])
    right_plot = cnpl.splineplot(cn._datahandler.validation_file_names[0],
                                label_quantiles = [0.2, 0.35, 0.5, 0.95],
                                channel = "Nd144Di",
                                x_scale = "linear",
                                y_scale = "linear",
                                ax = right_plot,
                                show = False)
    right_plot.set_title("Spline Plot:\nNd144Di", fontsize = 10)
    right_plot.set_xlim(right_plot.get_xlim()[0], right_plot.get_xlim()[1])
    right_plot.set_ylim(right_plot.get_ylim()[0], right_plot.get_ylim()[1]*1.1)
    right_plot.set_xlabel("original\nexpression\ndistribution")
    right_plot.set_ylabel("goal\nexpression\ndistribution")
    for ax in [left_plot, mid_plot, right_plot]:
        ax.tick_params(axis = "both", which = "both", labelsize = 8)
        ax.set_ylabel(ax.get_ylabel(), fontsize = 8)
        ax.set_xlabel(ax.get_xlabel(), fontsize = 8)

# %% [code]
def _generate_subfigure_d(fig: Figure,
                          ax: Axes,
                          gs: GridSpec,
                          subfigure_label: str):
    ax.axis("off")
    _figure_label(ax, subfigure_label, x = -0.1, y = 1.12)
    axes = gs.subgridspec(1,3, wspace = 1)
    
    from cytonormpy import Plotter
    cnpl = Plotter(cytonorm = cn)

    
    left_plot = fig.add_subplot(axes[0])
    left_plot = cnpl.emd(colorby = None, s = 10, edgecolor = "black", linewidth = 0.5, show = False, ax = left_plot)
    left_plot.set_xlabel("EMD after\nnormalization")
    left_plot.set_ylabel("EMD before\nnormalization")
    left_plot.set_title("EMD evaluation", fontsize = 10)
    
    right_plot = fig.add_subplot(axes[1])
    right_plot = cnpl.mad(colorby = None, s = 5, edgecolor = "black", linewidth = 0.3, show = False, ax = right_plot)
    right_plot.set_xlabel("MAD after\nnormalization")
    right_plot.set_ylabel("MAD before\nnormalization")
    right_plot.set_title("MAD evaluation", fontsize = 10)
    for ax in [left_plot, right_plot]:
        ax.tick_params(axis = "both", which = "both", labelsize = 8)
        ax.set_ylabel(ax.get_ylabel(), fontsize = 8)
        ax.set_xlabel(ax.get_xlabel(), fontsize = 8)

# %% [code]
fig = plt.figure(layout = "constrained",
                 figsize = (FIGURE_WIDTH_FULL, FIGURE_HEIGHT_FULL))

gs = GridSpec(ncols = 2,
              nrows = 4,
              height_ratios = [1,1,1.5,0.8],
              wspace = 0,
              hspace = 0.4)

a_coords = gs[0,:]
b_coords = gs[1,:]
c_coords = gs[2,:]
d_coords = gs[3,:]

fig_a = fig.add_subplot(a_coords)
fig_b = fig.add_subplot(b_coords)
fig_c = fig.add_subplot(c_coords)
fig_d = fig.add_subplot(d_coords)

_generate_subfigure_a(fig = fig,
                      ax = fig_a,
                      gs = a_coords,
                      subfigure_label = "A")

_generate_subfigure_b(fig = fig,
                      ax = fig_b,
                      gs = b_coords,
                      subfigure_label = "B")

_generate_subfigure_c(fig = fig,
                      ax = fig_c,
                      gs = c_coords,
                      subfigure_label = "C")

_generate_subfigure_d(fig = fig,
                      ax = fig_d,
                      gs = d_coords,
                      subfigure_label = "D")
plt.tight_layout()
plt.savefig("../figures/Figure_1.pdf", dpi = 300, bbox_inches = "tight")
plt.show()

# %% [code]
def _generate_subfigure_a(fig: Figure,
                          ax: Axes,
                          gs: GridSpec,
                          subfigure_label: str):
    DATASET_MAP = {
        "trussard": "Trussart",
        "dfci": "DFCI",
        "van_gassen": "Van Gassen"
    }

    ax.axis("off")
    _figure_label(ax, subfigure_label, x = -0.1, y = 1.1)
    axes = gs.subgridspec(1,2, wspace = 0, width_ratios = [1,0.5])
    data =  pd.read_csv("python_benchmark_functionwise.out")

    data.rename(columns={'clustering.1': 'clustering_time'}, inplace=True)
    data = data[data["clustering"] == True]
    data["dataset"] = data["dataset"].map(DATASET_MAP)
    
    data['validation_io'] = data['val_read'] + data['val_write']
    
    median_data = data.groupby('dataset').median().reset_index()
    
    columns_to_normalize = ['fcs_data_setup', 'clustering_time', 'quantile_calc', 'spline_calc', 'validation_io', 'transform'][::-1]
    median_data['total'] = median_data[columns_to_normalize].sum(axis=1)
    for col in columns_to_normalize:
        median_data[col] = median_data[col] / median_data['total']

    median_data.drop(columns=['total'], inplace=True)
    
    colors = plt.get_cmap('tab10')(range(len(columns_to_normalize))[::-1])
    barplot = fig.add_subplot(axes[0])
    median_data.set_index('dataset')[columns_to_normalize].plot(
        kind='bar', stacked=True, ax=barplot, color=colors)
    
    barplot.set_xticklabels(barplot.get_xticklabels(), rotation = 45, ha = "center", fontsize = 8)
    barplot.set_xlabel("")
    barplot.set_ylabel('Percentage of\nruntime', fontsize = 8)
    barplot.set_title('normalized time distribution per dataset', fontsize = 8)
    handles, labels = barplot.get_legend_handles_labels()
    print(labels)
    labels = [
        "Reference data setup", "Clustering", 
        "Quantile calculation", "Spline calculation",
        "Validation Files: I/O", "Normalization"][::-1]

    barplot.legend().remove()
    
    legend = fig.add_subplot(axes[1])
    legend.axis("off")
    legend.legend(handles[::-1], labels[::-1], title='Component', bbox_to_anchor=(0, 1), loc='upper left', fontsize = 8)


# %% [code]
def _generate_subfigure_b(fig: Figure,
                          ax: Axes,
                          gs: GridSpec,
                          subfigure_label: str):
    ax.axis("off")
    _figure_label(ax, subfigure_label, x = -0.1, y = 1.03)
    axes = gs.subgridspec(1,3, wspace = 0.6)

    top_plot = fig.add_subplot(axes[0])
    mid_plot = fig.add_subplot(axes[1])
    bottom_plot = fig.add_subplot(axes[2])
    
    df = pd.read_csv("van_gassen_unclustered.csv", index_col = [0])
    _comp_plot(df, top_plot, "Dataset 1: Van Gassen")

    df = pd.read_csv("dfci_unclustered.csv", index_col = [0])
    _comp_plot(df, mid_plot, "Dataset 2: DFCI")

    df = pd.read_csv("trussard_unclustered.csv", index_col = [0])
    _comp_plot(df, bottom_plot, "Dataset 3: Trussart")


# %% [code]
fig = plt.figure(layout = "constrained",
                 figsize = (FIGURE_WIDTH_FULL, FIGURE_HEIGHT_FULL/2))

gs = GridSpec(ncols = 2,
              nrows = 2,
              height_ratios = [1,1],
              wspace = 0,
              hspace = 0.8)

a_coords = gs[0,:]
b_coords = gs[1,:]

fig_a = fig.add_subplot(a_coords)
fig_b = fig.add_subplot(b_coords)

_generate_subfigure_a(fig = fig,
                      ax = fig_a,
                      gs = a_coords,
                      subfigure_label = "A")

_generate_subfigure_b(fig = fig,
                      ax = fig_b,
                      gs = b_coords,
                      subfigure_label = "B")


plt.tight_layout()
plt.savefig("../figures/Supplementary_Figure_2.pdf", dpi = 300, bbox_inches = "tight")
plt.show()
```
